# Effect of Geometric Sharpness on Translucent Material Perception

**DOI:** 10.1101/795294

**Authors:** Bei Xiao, Shuang Zhao, Ioannis Gkioulekas, Wenyan Bi, Kavita Bala

## Abstract

When judging optical properties of a translucent object, humans often look at sharp geometric features such as edges and thin parts. Analysis of the physics of light transport shows that these sharp geometries are necessary for scientific imaging systems to be able to accurately measure the underlying material optical properties. In this paper, we examine whether human perception of translucency is likewise affected by the presence of sharp geometry, by confounding our perceptual inferences about an object’s optical properties. We employ physically accurate simulations to create visual stimuli of translucent materials with varying shapes and optical properties under different illuminations. We then use these stimuli in psychophysical experiments, where human observers are asked to match an image of a target object by adjusting the material parameters of a match object with different geometric sharpness, lighting geometry, and 3D geometry. We find that the level of geometric sharpness significantly affects perceived translucency by the observers. These findings generalize across a few illuminations and object shapes. Our results suggest that the *perceived* translucency of an object depends on both the underlying material optical parameters and 3D shape. We also conduct analyses using computational metrics including (luminance-normalized) L2, structural similarity index (SSIM), and Michelson contrast. We find that these image metrics cannot predict perceptual results, suggesting low level image cues are not sufficient to explain our results.

## 1. Introduction

Most real-world materials are translucent, and humans interact with such materials often in their every day life. Examples include human skin, most foods, minerals, wooden objects, and chemical products (e.g., soap and wax). Compared to opaque materials, translucent ones have a characteristic appearance that results from light penetrating their surface, scattering internally, and eventually re-emerging from a different surface location. This kind of light-material interaction is known as *sub-surface scattering* (Ishimaru, 1991). The ubiquity and importance of translucent materials has long motivated research towards modeling and understanding their appearance. In recent years, there has been significant progress in developing rendering algorithms and representations that, by modeling the underlying physics, can create realistic reproductions of translucency (Jensen, Marschner, Levoy, & Hanrahan, 2001; Donner & Jensen, 2008; Gkioulekas et al., 2013). As with physics, there have likewise been previous efforts trying to decipher the other major aspect of translucent appearance, namely its *perception* by the human visual system (Fleming & Bülthoff, 2005; Motoyoshi, 2010; Anderson, 2011; Nagai et al., 2013; Gkioulekas et al., 2013; Xiao et al., 2014; Chowdhury, Marlow, & Kim, 2017; Marlow, Kim, & Anderson, 2017). Despite this, our understanding of translucency perception remains limited.

The study of the perception of translucency is confounded by the fact that there exists a tight coupling between how humans perceive the illumination, shape, and material properties of translucent objects (Xiao et al., 2014; Marlow et al., 2017). Recently, Chowdhury et al. (Chowdhury et al., 2017) showed that the perception of the 3D shape of a translucent object is different from that of an opaque one. They created stimuli by manipulating the spatial frequency of relief surfaces for both translucent and opaque objects. Using these stimuli, they found that observers judged translucent objects to have fewer bumps than opaque objects with the same 3D shape. Additionally, they observed that the perceived local curvature was underestimated for translucent objects relative to opaque objects.

Motivated by these intriguing findings, in this paper we continue the investigation of the interplay between the perception of 3D shape and material for translucent objects. We focus on a specific type of geometric features, namely thin geometric structures such as edges and depth discontinuities. We will use the term *geometric sharpness* to refer broadly to the presence of such features. Our focus on geometric sharpness is justified from previous studies in both physics (Gkioulekas, Walter, Adelson, Bala, & Zickler, 2015; Zhao, Ramamoorthi, & Bala, 2014) and perception (Fleming & Bülthoff, 2005; Gkioulekas et al., 2013), indicating that the presence of thin geometric features is critical for discriminating between translucent materials. In particular, we investigate the following questions:

- How does geometric sharpness affect the perception of translucent material properties?
- Can we manipulate geometric sharpness to make an object appear more or less translucent?

To answer these questions, we first review related previous work in Section 2. Then, we start our investigations by synthesizing a large number of image stimuli, with greatly varying 3D shapes, illumination, and optical properties, detailed in Section 3. We use these stimuli to conduct psychophysical studies, the results of which are statistically analyzed in Section 4. Finally, in Section 5, we discuss implications of our experimental results. Specifically, our findings indicate that 3D geometric sharpness affects translucent material perception in such a way that blurring geometric details cause objects with shallow relief to appear more translucent than the objects with sharp geometry.

## 2. Previous work

Illumination, material properties, and 3D shape all contribute to the retinal 2D image that humans perceive. The human visual system subsequently uses these images, among other things, to infer material properties of the objects contained in them. In this process, the visual system takes into account the effects of the ancillary shape and illumination properties (Barron & Malik, 2015).

Previous studies on the perception of opaque objects have demonstrated that the shape of an object can affect its perceived material properties (Ho, Landy, & Maloney, 2008; Todd & Mingolla, 1983; Vangorp, Laurijssen, & Dutré, 2007; Wijntjes & Pont, 2010; Marlow & Anderson, 2013, 2015; J. Kim, Marlow, & Anderson, 2012). By varying the surface geometry, it is possible to influence how glossy an object is perceived by human observers to a significant extent. For example, M. Kim, Wilcox, and Murray show that perceived three-dimensional shape plays a decisive role in perception of surface glow. Specifically, by manipulating cues to 3D shape while holding other image features constant, the perception of glow can be toggled on and off.

Conversely, the perception of 3D shape is affected by the surface’s optical (i.e, reflectance) properties. For example, it has been shown that the same surface can be perceived to have different 3D geometry when its reflectance becomes more or less specular (Doerschner, Yilmaz, Kucukoglu, & Fleming, 2013; Adams & Elder, 2014; Sawayama & Nishida, 2018). This coupling between the perception of 3D shape and surface reflectance properties has also been demonstrated in objects with non-specular reflectance (Todd, Egan, & Phillips, 2014). To mention one example, human observers perceive velvet materials as flatter than matte-painted materials of the same shape (Wijntjes, Doerschner, Kucukoglu, & Pont, 2012). Illumination also enters this interplay, as previous findings show that changes in the lighting of an object can strongly affect its perceived shape (Norman, Todd, & Orban, 2004) and reflectance (Olkkonen & Brainard, 2011; Marlow, Kim, & Anderson, 2012).

Similar to human perception of opaque objects, it has been shown that humans also use a variety of image cues to detect subtle differences between translucent objects, as well as to make qualitative inferences about their optical parameters (Fleming, Jensen, & Bülthoff, 2004; Fleming & Bülthoff, 2005; Motoyoshi, 2010; Anderson, 2011; Nagai et al., 2013; Gkioulekas et al., 2013). These findings suggest that the perception of translucency is influenced by surface reflectance (gloss versus matte) (Motoyoshi, 2010), illumination conditions (smooth versus harsh) (Xiao et al., 2014), as well as surface geometry (Chowdhury et al., 2017; Marlow et al., 2017). Additionally, it has been suggested that the coupling between the perception of shape and material properties is more complicated for translucent objects than for opaque ones (Anderson, 2011). From the perspective of physics, it has been shown that estimating 3D shape from translucent objects is challenging (Koenderink & Doorn, 2001; Inoshita, Mukaigawa, Matsushita, & Yagi, 2012) and a recent study found that recovered surface normals of shallow relief objects are smoother than opaque objects with the same 3D shape, suggesting volumetric scattering has a blurring effect on the photometric normals (Moore & Peers, 2013). This finding partially inspired our work to study whether similar interaction between 3D shape and scattering parameters also exists in perception.

In a recent work, Chowdhury et al. used rendered stimuli that were either opaque or translucent to study the effects of translucent material properties on perception of 3D shape. In this paper, we study how 3D shape affects the perception of translucency by systematically varying the geometric sharpness and optical density of relief objects. Thus, instead of being opaque or translucent, our stimuli vary in degree of translucency. Our focus on sharp geometric features is motivated by prior research on the physics of light scattering, which shows that imaging systems can only accurately measure scattering parameters in the presence of such features (Zhao et al., 2014); and that intensity profiles across geometric edges of translucent objects provide rich information about the object’s scattering material (Gkioulekas et al., 2015). Specifically, we design a matching experiment to measure the correlation between geometric sharpness and perceived material translucency. By analyzing the data collected through this experiment, we demonstrate that geometric sharpness indeed affects the perception of translucency for objects with lower reliefs. Specifically, smoother geometry usually leads to higher levels of perceived translucency. Stated differently, when shown two objects with identical optical properties, humans tend to perceive the one with smoother geometric structures as more translucent.

## 3. General Methods

### Stimuli

We generated our visual stimuli using 3D models representing a cube with the characters “target” or “match” raised above the top surface. In our experiments, we focus on the effect of the geometry of the relief (instead of the cube edges and corners) and only show the top surface. Figure 7 shows examples of the resulting rendered images.

The stimuli were rendered using physically accurate simulation of subsurface scattering, implemented by the Mitsuba renderer (Jakob, 2013). In the following, we provide details about the geometric, illumination, and material models we use for stimuli generation.

### Geometric

Figure 3 illustrates the 3D geometry of our rendered objects. In our asymmetric matching experiment, we rendered different letters for the surface relief for match and target. Besides using different letters, the base (top) surfaces are also slightly different. Here, we use “target” stimuli as examples to illustrate image examples. The object was modeled as a cube that sits on a flat surface (see figure 4). We used two-dimensional height fields, applied at the top surface of a cube (size 128*mm* × 96*mm* × 30*mm*), to control the shape of our relief models. The height fields were represented as z(x,y), and by applying low-pass (e.g., Gaussian) filtering to the height field, we could easily manipulate the object’s geometric sharpness. The relief heights were ranged from 0.5*mm* to 2.5*mm*. The kernel standard deviation was ranged from 0.08*mm* to 0.56*mm*. We used two types of relief as our stimuli in Experiment 1 and Experiment 2, respectively. The positive relief had the pattern raised above the top ground surface and negative relief had the pattern below sunken into the ground surface (Figure 6 and figure 10 for examples).

**Figure 1:**
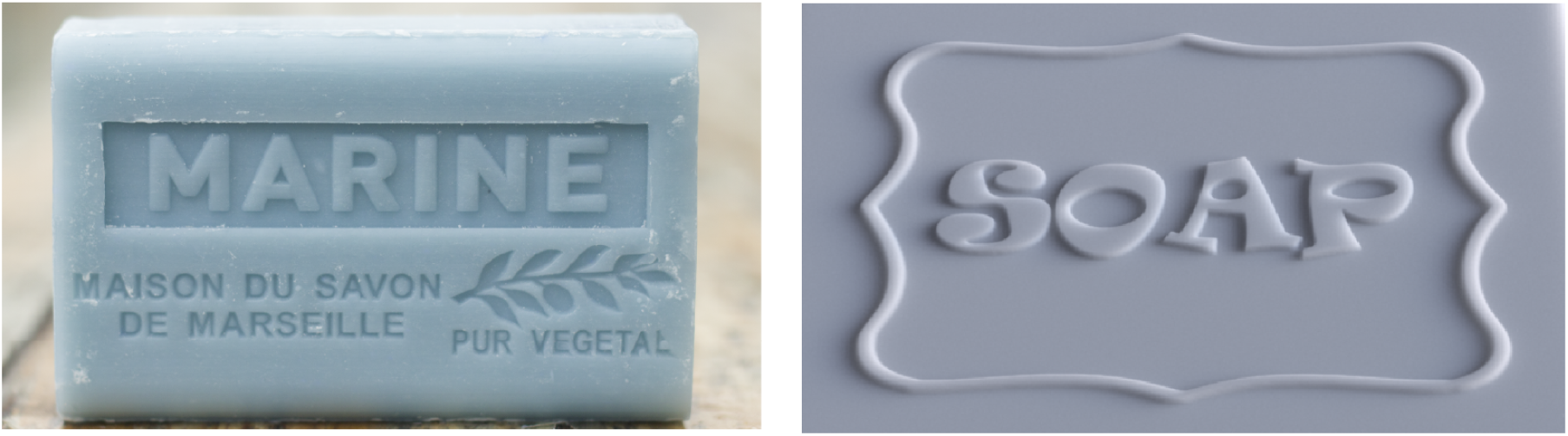
**A.** Photograph of a translucent soap bar with surface reliefs. Previous work indicates that the sharp edges of the translucent objects provide rich physical information about their material properties. Our findings indicate that sharp edges are also an important cue for how these material properties are perceived by humans. **B.** Inspired by this photograph, we render synthetic translucent images with a similar relief geometry, and use similar images as stimuli for psychophysical experiments.

**Figure 2:**
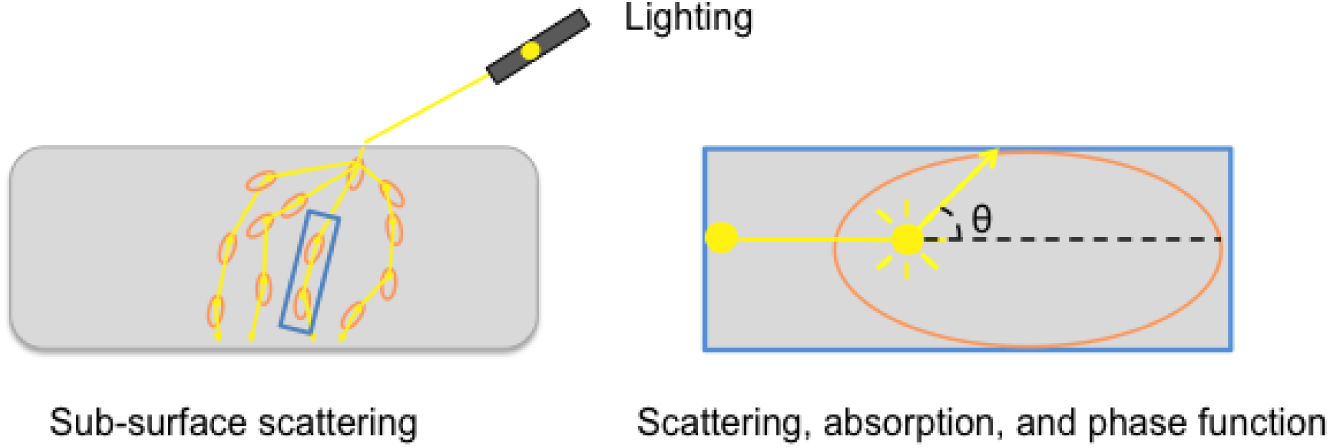
Illustration of the physics of subsurface scattering using the radiative transfer framework. Left: Translucent appearance is caused by light scattering inside the volume of an object, which can be described by the radiative transfer framework (Chandrasekhar, 1960). Right: A closer look of subsurface scattering: Between scattering events, light travels for distances determined by the object’s extinction coefficient. At each scattering event, light is either absorbed or scattered towards different directions, depending on the object’s volumetric albedo. Finally, when light is scattered, the phase function describes the angular distribution to each new direction of travel.

**Figure 3:**
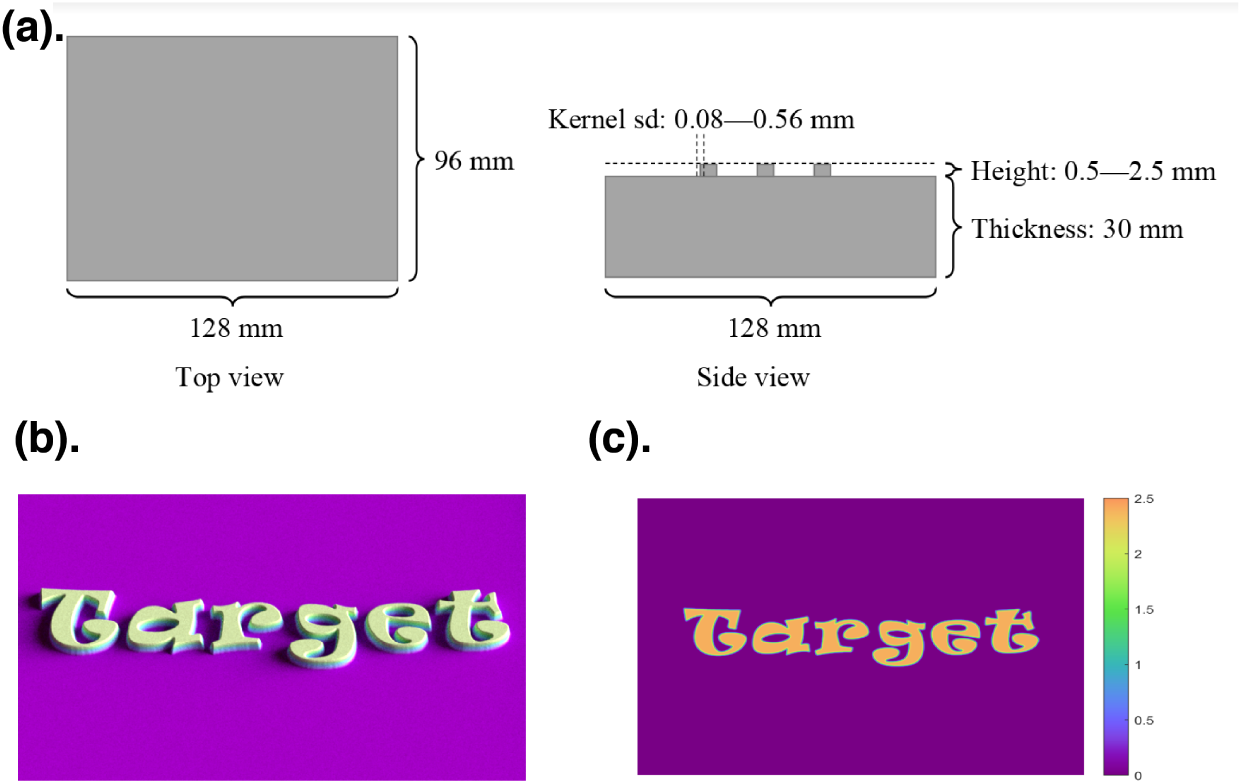
Illustration of the 3D geometry of the rendered objects. (a) Dimensions of the object. (b). 3D model of TARGET object. (c). Height maps of the surface relief. Different colors represent different height (z axis) at each pixel locations.

**Figure 4:**
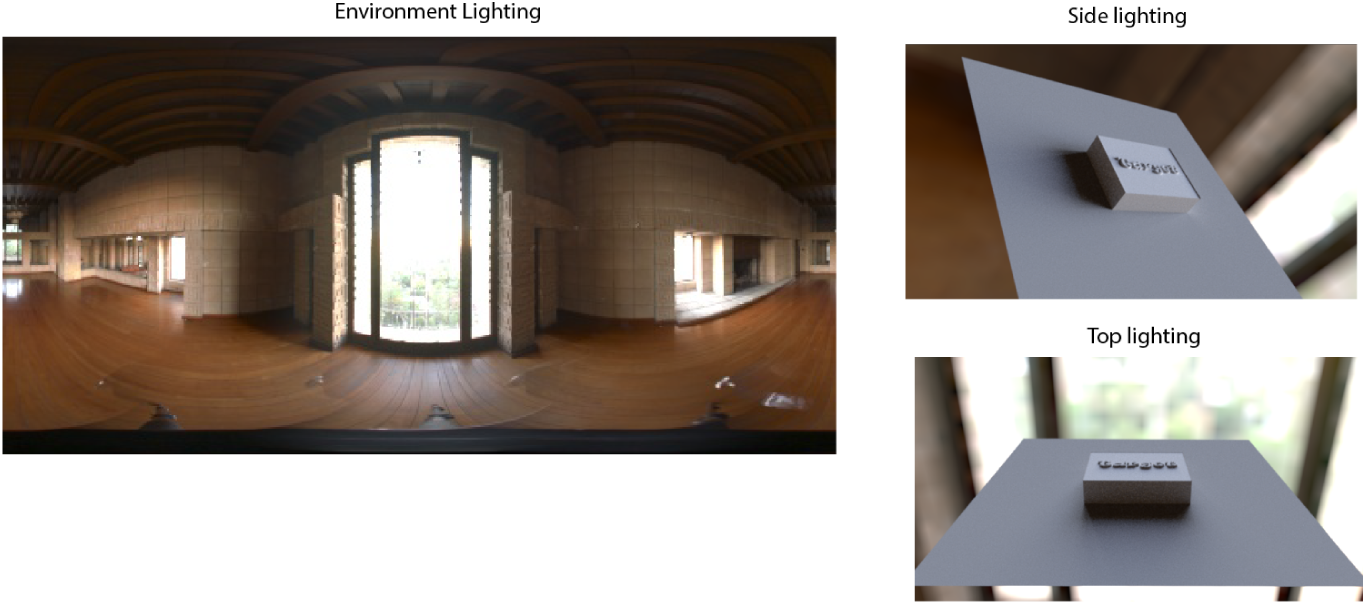
Left: The “Ennis” environment map used to generate our stimuli. Right: Example renderings of the stimuli object under two lighting conditions. In the “side-lighting” condition, the tall floor window is to the right of the object. But there are also light sources (smaller windows) on the left. In the “top lighting” condition, the same tall floor window that provides most direct lighting is behind the object. The camera views of the example renderings are for illustration purpose ONLY and are slightly DIFFERENT from what we used in the paper.

### Material

We used the radiative transfer framework (Chandrasekhar, 1960) to describe the scattering of light inside translucent objects. Radiative transfer is widely used in applied physics, biomedicine, remote sensing, and computer graphics. Under this framework, the optical properties of a homogeneous translucent material are described using three parameters (see Figure 2):

- The *optical density* is a measure of physical translucency. Also known as the extinction or total attenuation coefficient, it specifies how frequently light scatters within the material. An optical density of X mm^−1^ means light on average travels (in straight lines) for 1/X mm before being scattered by the medium. Thus, 1/X is usually termed as the “mean free path”. A higher density, or a lower mean free path, means more frequent scatterings of light. Optically thin materials (i.e. those with low optical densities) are semi-transparent. Optically thick materials, on the other hand, can appear mostly opaque, as light cannot travel very far inside the material due to very strong attenuation.
- The *volumetric albedo* determines how much light is absorbed and how much light is scattered to other directions, every time a scattering event takes place.
- The *phase function* determines how scattered light is distributed to different directions when scattering events take place.

### Lighting

To provide natural illumination, we used environmental lighting for our renderings. Specifically, we used the publicly available “Ennis” lighting environment (see Figure 4) that captures the illumination of the dining room of the Ennis-Brown house in Los Angeles, California (Debevec, 1998). Since this lighting condition is highly directional due to the bright glass door, we rotated it to create both top lighting and side lighting. The right panels in Figure 4 illustrated the two lighting conditions we used in the experiments with two example renderings. In the “side lighting” condition, the tall glass door is at the right side of the object while in the “top-lighting” condition, the tall glass window is behind the object. Since the viewer would look at the object mostly from above, we call this lighting condition “top lighting”. One can tell the lighting direction by looking at the cast shadows. Our choice of environmental lighting is due to the observation that materials rendered under natural lighting appear more realistic than materials rendered under directional lighting.

In order to ensure that our matching experiments is asymmetric, we used different extruding characters (geometry) and lighting geometry to render the “target” and “match” objects. The lighting directions for both side and top lighting slightly differ from the target and match images (see Figure 5). Specifically, we rotated the environmental map slightly (15 degree) in the scene depending on whether it is “target” or “match” to avoid exactly identical illumination conditions.

**Figure 5:**
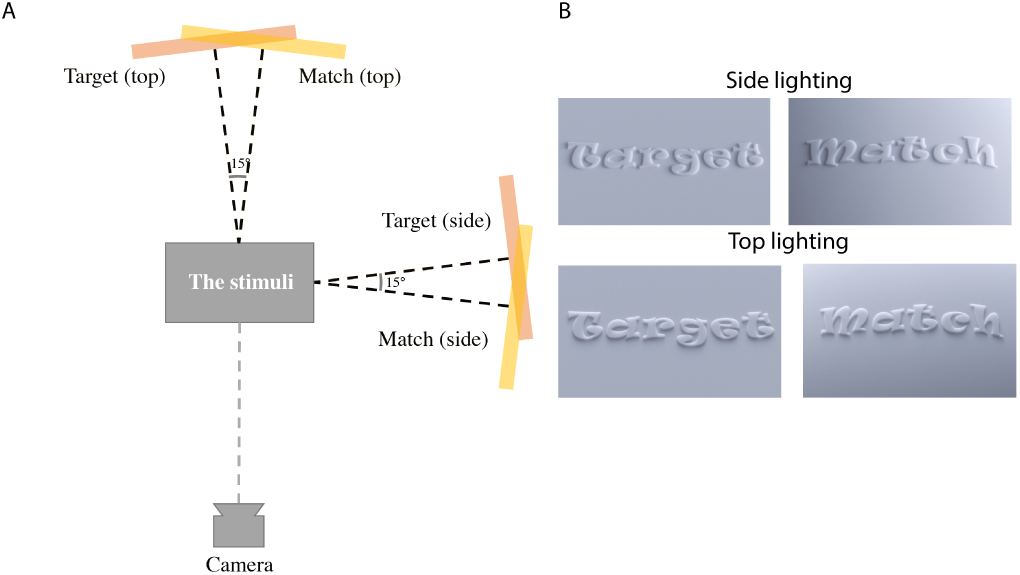
A. Top-view illustration of our virtual setup in for rendering the stimuli. The light pink and yellow rectangles represent the light source. B. To make sure our experiment is asymmetric, we used different relief letter and slightly different lighting to illuminate “target” and “match” images. Examples of “target” and “match” relief images used in Experiment 1 for relief height =0.5*mm* and blur level 0.2*mm*.

### Sampling of model parameters

We varied the geometric, material, and lighting properties of the stimuli (see example images in Figure 6).

For geometry, we used reliefs with five heights (0.5, 1.0, 1.5, 2.0, 2.5*mm*) where smaller values indicate lower reliefs. For each height, we apply Gaussian blurring to the underlying height fields using four kernel standard deviations (0.08, 0.2, 0.4, and 0.56 *mm*). Larger blurring kernels result in stimuli with lower geometric sharpness. The thickness of the rendered object is 30*mm*.

**Figure 6:**
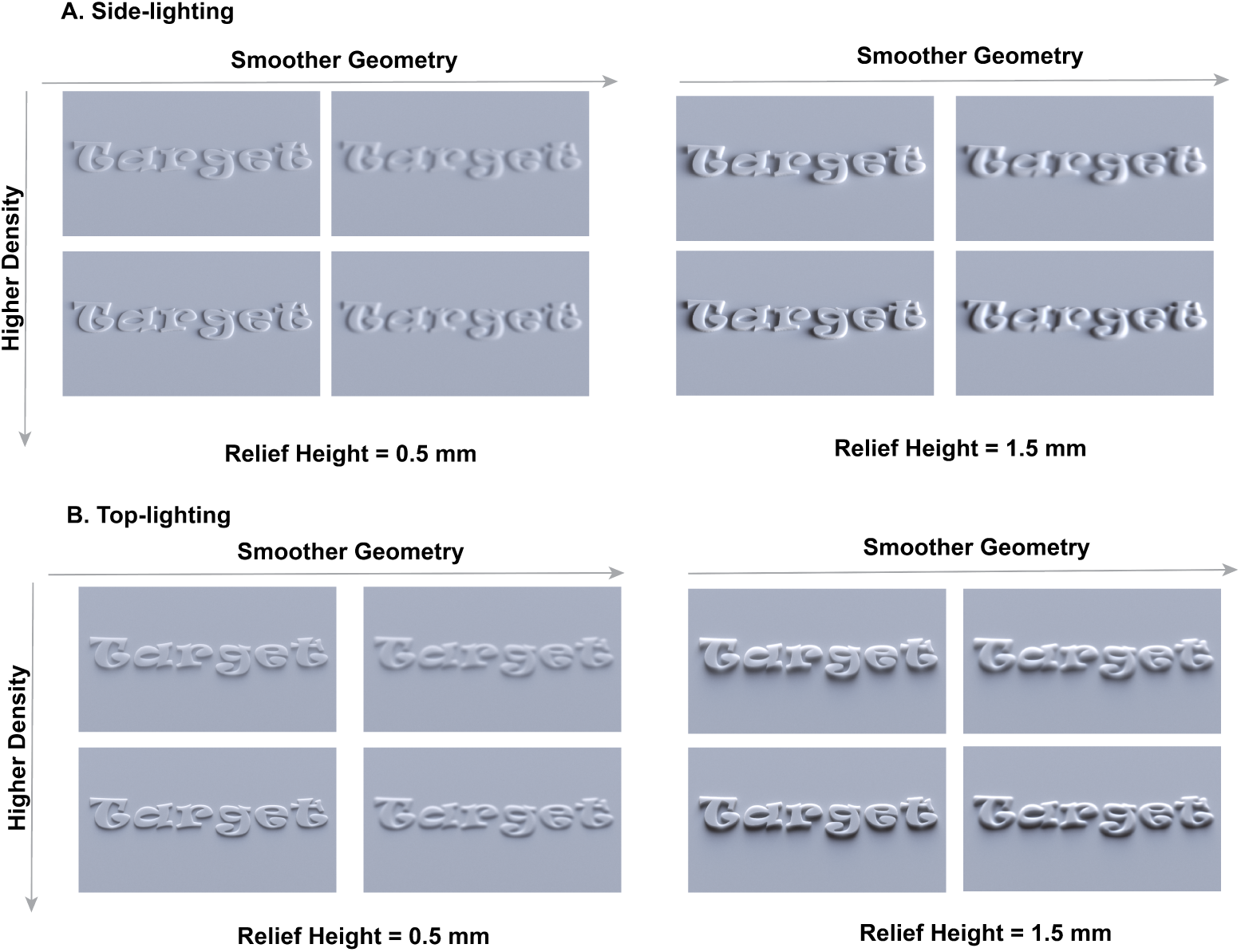
Experiment Stimuli. We manipulated the 3D shape of our stimuli by adjusting the extrusion height and geometric sharpness. For each shape, we render the object with varying optical densities and illumination conditions. Here, we only showed “target” image as examples.

For material scattering parameters, we used 15 optical density values sampled logarithmically between 0.7 and 3 (*mm*^−1^). We fix volumetric albedo to 0.9, and the phase function to be uniform.

Finally, we render the objects using both side and top environmental lighting. In total, our stimuli dataset consists of 4 (densities) × 5 (heights) × 4 (blurs) × 2 (lightings) = 160 target images. For each target image, we also rendered 15 (densities) × 1 (height) × 1(blur) × 1 (lighting) = 15 match images for a different but fixed blur level (with kernel size 0.24*mm*). In total we rendered 2400 images.^1^

The rendering, lighting and experimental procedure for Experiment 2 (negative relief) is the same as above except for the geometry of the stimuli.

### Procedure

We used the asymmetric matching method to obtain psychophysical measures of the perceived translucency (Brainard & Wandell, 1992). Previously, asymmetric matching was used in measuring color constancy where subjects set asymmetric color matches between a standard object and a test object that were rendered under illuminants with different spectral power distributions. Besides color vision, asymmetric matching has also been used in material perception such as effect of surface gloss on color perception (Xiao & Brainard, 2008), effect of illumination on perception of lightness and glossiness (Olkkonen & Brainard, 2010), effect lighting direction on translucency (Xiao et al., 2014), effect of distortion on transparency (Fleming, Jäkel, & Maloney, 2011), effects of optical properties on perception of liquids (Assen & Fleming, 2016), etc. It is an important method to measure how a particular factor affects material perception by preventing observers from performing low-level image matching.

To this end, we created a browser-based experiment interface to collect the matching data (see Figure 7). Even though the experimental interface was coded using JavaScript (instead of MATLAB or PsychoPy), we didn’t collect the data on Mechanical Turk but simply used the browser as an interface to collect the responses from the observers. The interface displayed two images. The object shown in the left (target) image has a fixed optical density. The interface provided a slider that allows observers to change the optical density of the object shown in right (match) image.

**Figure 7:**
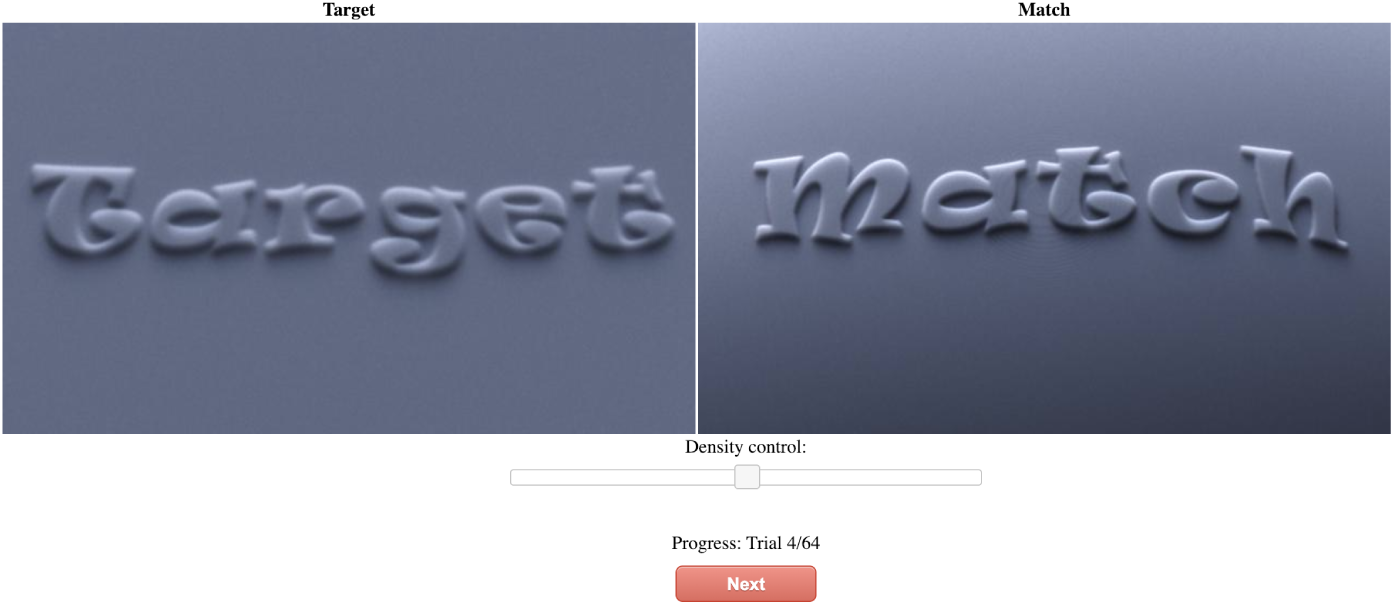
Experimental user interface of the translucency matching experiment. Human observers are asked to use the slider to adjust the optical density of the right (match) object, until its perceived material properties matches the appearance of the left (target) object. The match image always have a fixed blur level within each trial, which is different from that of the target image.

Before the start of the experiment, observers were told that the images they would see were objects made of translucent materials, and they were shown some example stimuli images. As the experiment started, observers were shown multiple pairs of images using the experiment interface. For each image pair, the specific instructions that were given to the observer were the following. The reliefs in the images you will see in different trials are made with different materials. Your task is to match the perceived material properties of the soap in the right image to that in the left. To do so, you adjust the “opacity” of the object in the right image (match) to match the perceived translucency to the object on the left (target). Notice that from trial to trial, the lighting direction can change (either side-lit or top-lit).

Once they were satisfied with the match, the observers pressed the “next” button to confirm their response. Following a response, the screen was blanked for 2 seconds, and subsequently a new image pair was shown. We call each such matching process a trial.

At each trial, the match object and target object were rendered with the same extrusion heights and the general similar lighting directions (both side or top but slightly different, see Lighting section). The target object had a specific density value and a specific amount of blur, whereas the match object always has a fixed blur level (with kernel sd of “0.24 mm”), which was never the same as the target. In total, the experiment consisted of 160 trials, divided into 4 blocks of 40 trials each. There was no time limit for finishing each trial, and observers could take a break between the blocks.

In Experiment 2, we used the same procedure as described above for a different group of observers (see section Observers).

### Display

Observers performed the experiments in a dark room and the images were displayed on a 27-inch iMacPro monitor (Apple, Inc., Cupertino, CA; dynamic range: 60:1; color profile: sRGB; maximum luminance: 340 cd/m). The height of the image was 80 mm and the observers sat approximately 50 cm away from the monitor. The stimulus subtended 8.6 degrees in visual angle.

Our rendered “RAW” images were float-valued and contained radiometric quantities (i.e., radiance values). To properly display the RAW images, we convert them into the sRGB color space using the standard approach. This conversion is similar to applying *I*_*sRGB*_ = *I*_*Raw*_ (1*/*2.2), although they are not strictly identical. This conversion of color space (from linear RGB to sRGB) is the standard way to display rendered images (or measured RAW photos if that matters). Our monitors use the sRGB color gamut and are properly calibrated. This means, when displaying a sRGB image, light physically emitted by our monitors will closely resemble the radiometric quantities recorded by the original simulated results.

## 4. Experiment 1: Positive Relief

### Introduction

In Experiment 1, we measured the effects of geometric sharpness on perceived translucency with the afromentioned asymmetric matching procedure using positive relief objects. Throughout the conditions, we also varied optical density, relief heights, lighting conditions.

### Observers

13 observers (8 women; with mean age 22 years, and standard deviation of age 2.5 years) participated in this experiment. All observers reported normal visual acuity and color vision. All observers participated for credit in an introductory psychology course. The procedures were conducted in accordance with the Declaration of Helsinki and were approved by the Human Research Ethics Advisory Panel at American University.

### Results and Discussion

Figure 8A demonstrates the mean match density across 13 observers versus blur levels for condition (height=1.0*mm* and lighting= “side-lit”), which is the same as the second panel on the top raw in figure 9. We make the following observations.

**Figure 8:**
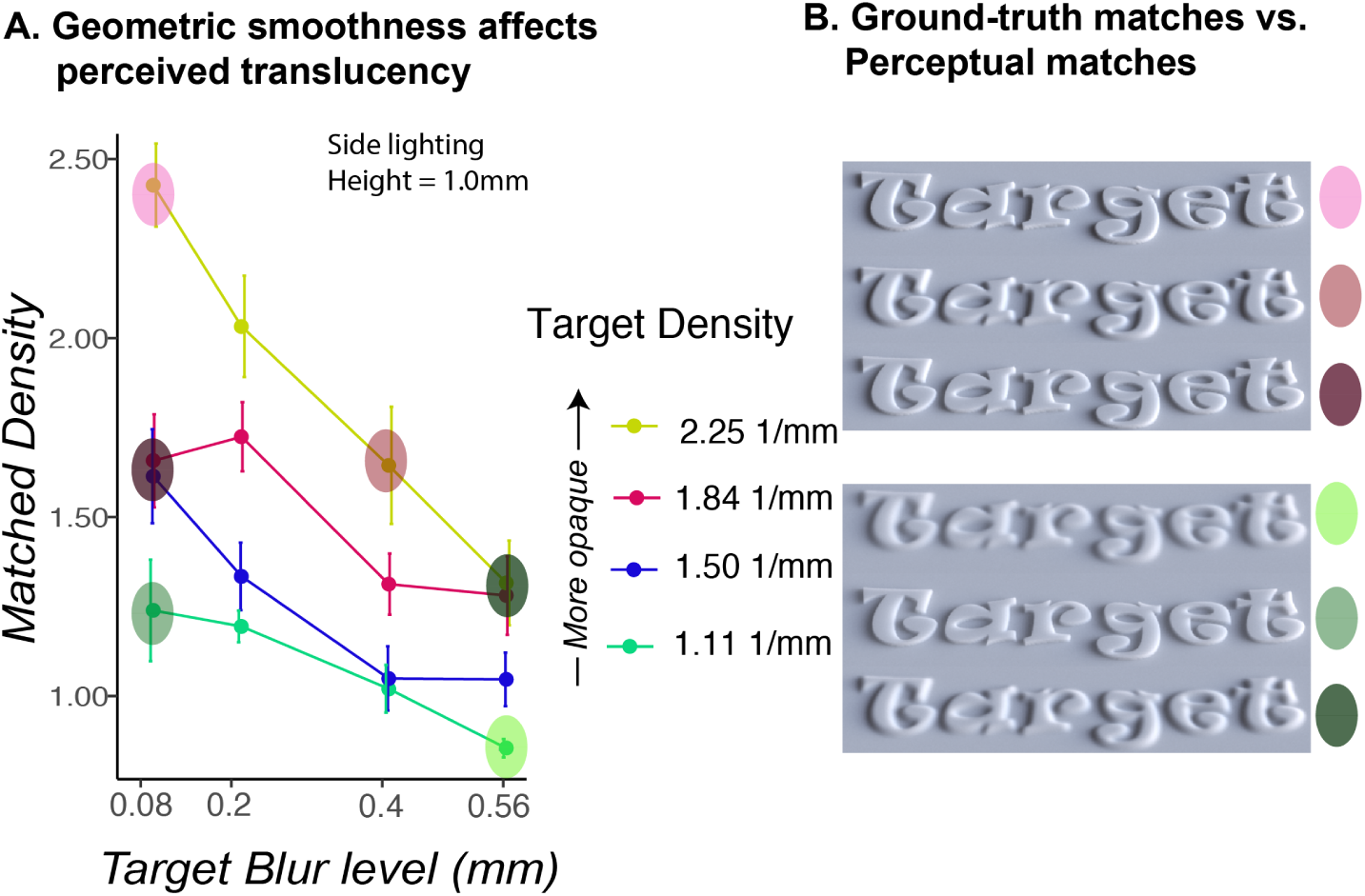
Asymmetric matching results and image demonstrations for Experiment 1 with relief height 1.0*mm*: blurring (smoothing geometric details) cause the object to appear more translucent (less opaque). **A.** Mean match density across observers versus the level of blur. Different colors represent stimuli with different optical densities. **B.** Demos of perceptually equivalent image pairs versus pairs that have the same physical densities. Higher values indicates more opaque appearance. Top panel: observers perceive a target object with higher density (middle image, light maroon dot) and smooth geometry to be equally translucent to an object with lower density and sharp geometry (lower image, dark maroon dot). The top image (pink dot) shows the physical ground-truth image of the object with the same density as the target. Lower panel: observers perceive a target object with lower density (middle image, green dot) with sharp geometry to be equivalent to an object with higher density with smooth geometry (lower image, dark green dot). The top image shows the physical ground-truth image of the object with the same density as the target (lime green dot).

**Figure 9:**
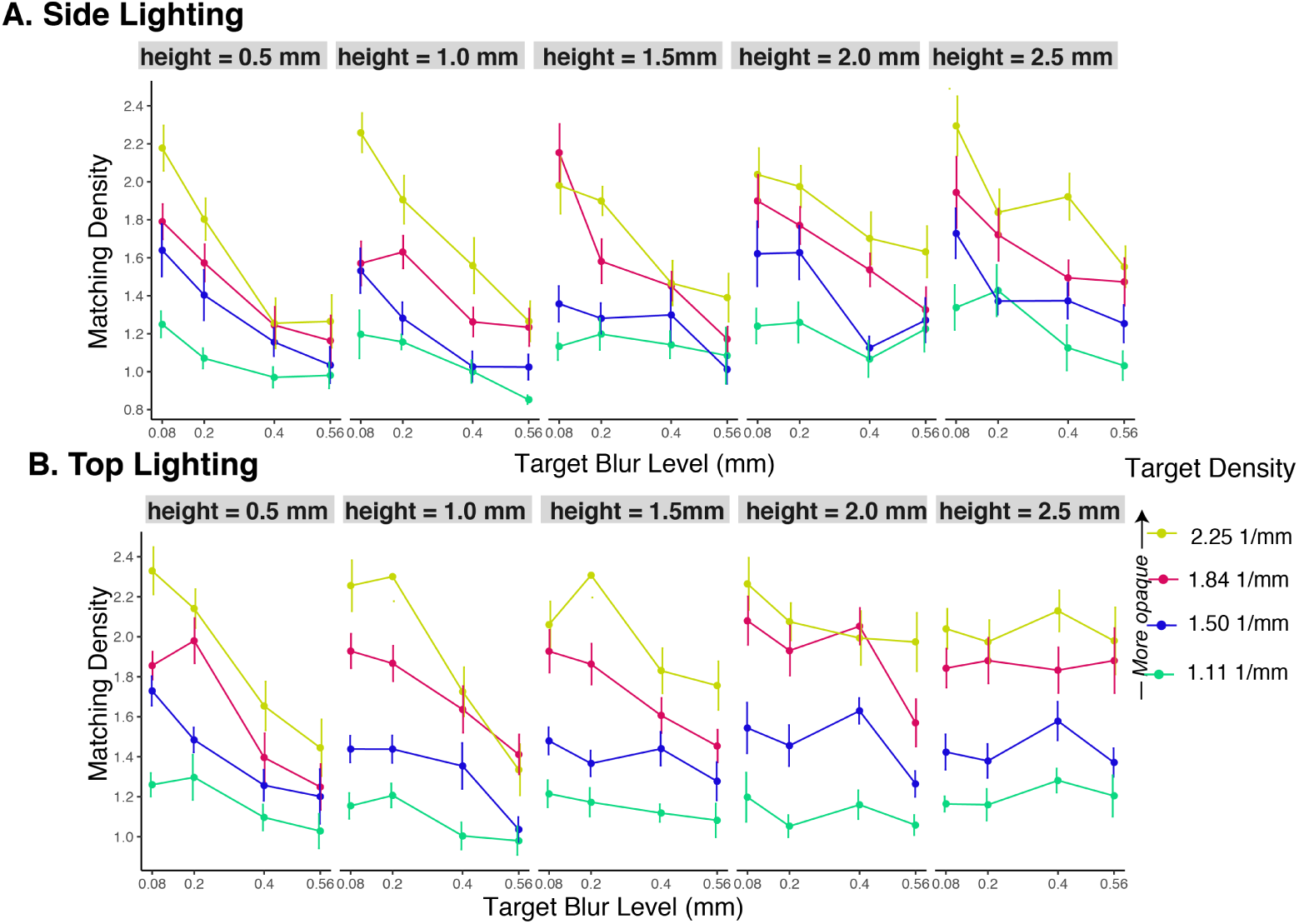
Results from Experiment 1 (Positive relief) of all relief heights and lightings. **A.** Effect of geometric smoothness (applying Gaussian blur to 2D height fields) on the matching results for all relief heights under side lighting. Each panel plots the level of blur versus averaged matched density for a specific relief height value. Higher value on y-axis represents more opaque materials and lower value represents more translucent materials. From the left panel to the right, the relief heights increase. Different color represents different optical densities for the target as shown in the figure legend. **B.** Same plots as shown in **A** but for top-lighting conditions. For both lighting conditions, when the relief heights not at the highest level (left four panels), as blur level increases, mean matched density decrease, suggesting observers perceive geometric smoothed relief objects to be more translucent. At the highest relief level, for top lighting the effect is diminished (resulting in nearly flat lines).

We first note that observers can distinguish different optical densities when the target and the match have the same height and blur values. For these conditions, as the material optical density of the target increases, the matched density increases as well (e.g., the yellow line is well above green line in Figure 8A). This shows observers perceive objects with higher optical density to be more opaque.

On the other hand, as shown in Figure 8A, when the blur level increases, the observers’ matched density decreases for each fixed target density. Put another way, **to match the translucent appearance of the target object with the higher blur level, observers need to decrease the density of the match object.** Equivalently, this suggests that smoothing the height fields makes the object appear less dense optically (more translucent). To visually demonstrate this result, Figure 8B shows representative triplets of images, where the image of the target object is compared with the image of the perceptual match, as well as the image of the object that has the same ground-truth density as the target. The top panel of Figure 8B shows the effect of blur on translucent appearance of selected images. The middle image of the top panel is a cross section of a blurred object which corresponds to the maroon dot on the data plot in Figure 8A. The bottom image of the panel shows the image rendered with mean matched density across observers (dark maroon). Even though the perceptually matched object in this image has lower density and sharper features (lower blur level) than the target (red dot on data plot in Figure 8A), observers perceive them to be similar in translucency. In contrast, the top image shows the object rendered with the same physical density as the target but with a lower blur level (pink dot on the data plot). Observers perceive this image to be more opaque than the target.

The bottom panel illustrates another example of the effect of blurring, where a translucent object with sharper features (middle, green dot) is perceived to be equivalent to a smoothed object (higher blur level) with higher density (bottom, dark green dot) in contrast with the object with the same physical density (top, lime dot). Together, this demonstrates that sharp geometries affect translucent appearance in such a way that a geometrically smoothed object appears more translucent than the sharp object that has the same optical density.

We further examine results for all other experimental conditions. Figure 9 shows the average matching results for the relief objects for all five height conditions under both side (top panel) and top lighting (bottom panel) conditions. The format of Figure 9 is the same as Figure 8A where the x-axis represents the level of blurring applied to the height fields and the y-axis represents the matched density. From left to right, the plots show results for conditions with increasing heights of the relief. The top panel in Figure 9 shows that geometric smoothness has a significant effect on perceived translucency, meaning the mean matched density decreases as the blur increases. The prominence of this effect depends on the optical density. Blurring has a stronger effect for the high-density conditions (yellow, blue, red lines) than the low-density (green lines). The same trend is observed for the top lighting conditions (bottom panel in Figure 9). Blur has a strong effect on low relief heights (left two plots) and the effect of blur on matched density is flattened for the higher relief values (right most panel).

We perform a three-way within-subject ANOVA on the difference between matched density and the target density with blur level, relief extrusion height and the lighting direction as independent variables. To summarize, we find a significant effect of blur on perceived translucency considering all the conditions such that as blur increases, observers perceive the target objects to be more translucent. We also find significant main effects of relief heights and lighting on perceived translucency.

The results are as follows.

- *Blur level* has a significant main effect (*F* (3, 2000) = 67.985, *p <* 0.001), such that as the amount of blurring increases, the value of *d*_match_ − *d*_density_ decreases. The value is negative and its magnitude ∥*d*_match_ − *d*_density_∥ becomes larger, indicating that blur has a stronger effect for targets with higher densities.
- *Relief height* has a significant main effect (*F* (4, 2000) = 5.497, *p <* 0.001), such that larger values of relief height make the objects be perceived as less translucent (higher values of *d*_match_ − *d*_density_). There is no significant interaction between blur level and relief height (*F* (12, 2000) = 1.701, *p* = 0.0605).
- *Lighting direction* also has a significant main effect (*F* (1, 2000) = 22.788, *p <* 0.001). There is no significant interaction between lighting and blur level (*F* (3, 2000) = 1.122, *p* = 0.095).

## 5: Experiment 2: Negative Relief

### Introduction

The relief of objects in the real world (e.g., bas-relief sculptures or soaps) can be extruded positively or negatively. To discover whether the effect of 3D shape on perceived translucency we observed in Experiment 1 can be generalized to other shape, we rendered a set of similar stimuli with negatively extruded geometries and measured the effects of geometric sharpness on perceived translucency.

### Observers

Another eleven observers (8 women; mean age = 25 years, *SD* = 3.5 years) participated in the experiment where we used negatively extruded objects (Experiment 2). All the other aspects of the procedure and methods are the same as Experiment 1.

### Results and Discussion

Figure 10 shows example stimuli with negative relief under two lighting conditions (side lighting, top lighting) and two relief heights (*height* = 0.5, 1.5*mm*). Within each panel, the stimuli have increasing optical densities from top to bottom rows (*d*_target_ = 1.11, 2.25(1*/mm*)) and the blur level is increased from left to right (*blur* = 0.08, 0.56*mm*). All the other rendering parameters are identical to the stimuli used in Experiment 1.

**Figure 10:**
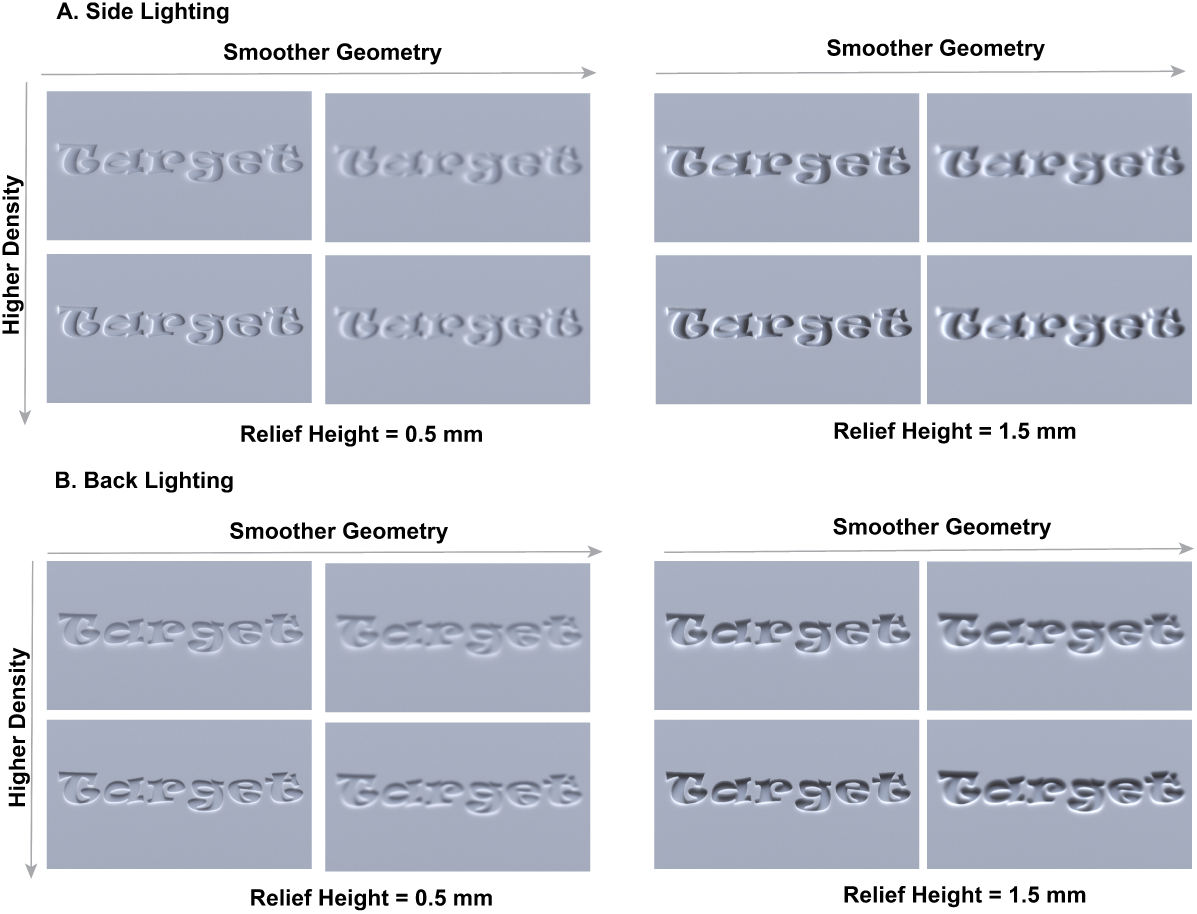
The conditions used in Experiment 2 are the same as in Experiment 1, except that the shape of the objects is different (negatively relief). **A.** From left to right, we applied increasing Gaussian blur to the height fields and from top to bottom, the object has increasing optical density. **B**. Same as A but with higher relief. **C**. Same as A but under top lighting. **D**. Same as B but under top lighting.

Figure 11 shows an example of the matching results for Experiment 2 with negative relief (*height* = 0.5*mm, lighting* = “*top*”), which is the the same figure in second column of the lower row in figure 12. Similar to Figure 8, Figure 11 plots the matched densities against different levels of blurring for each relief height level under top lighting. We show image examples of how blur affect perceived translucency, where the image of the target object is compared with the image of the perceptual match, as well as the image of the object that has the same ground-truth density as the target.

**Figure 11:**
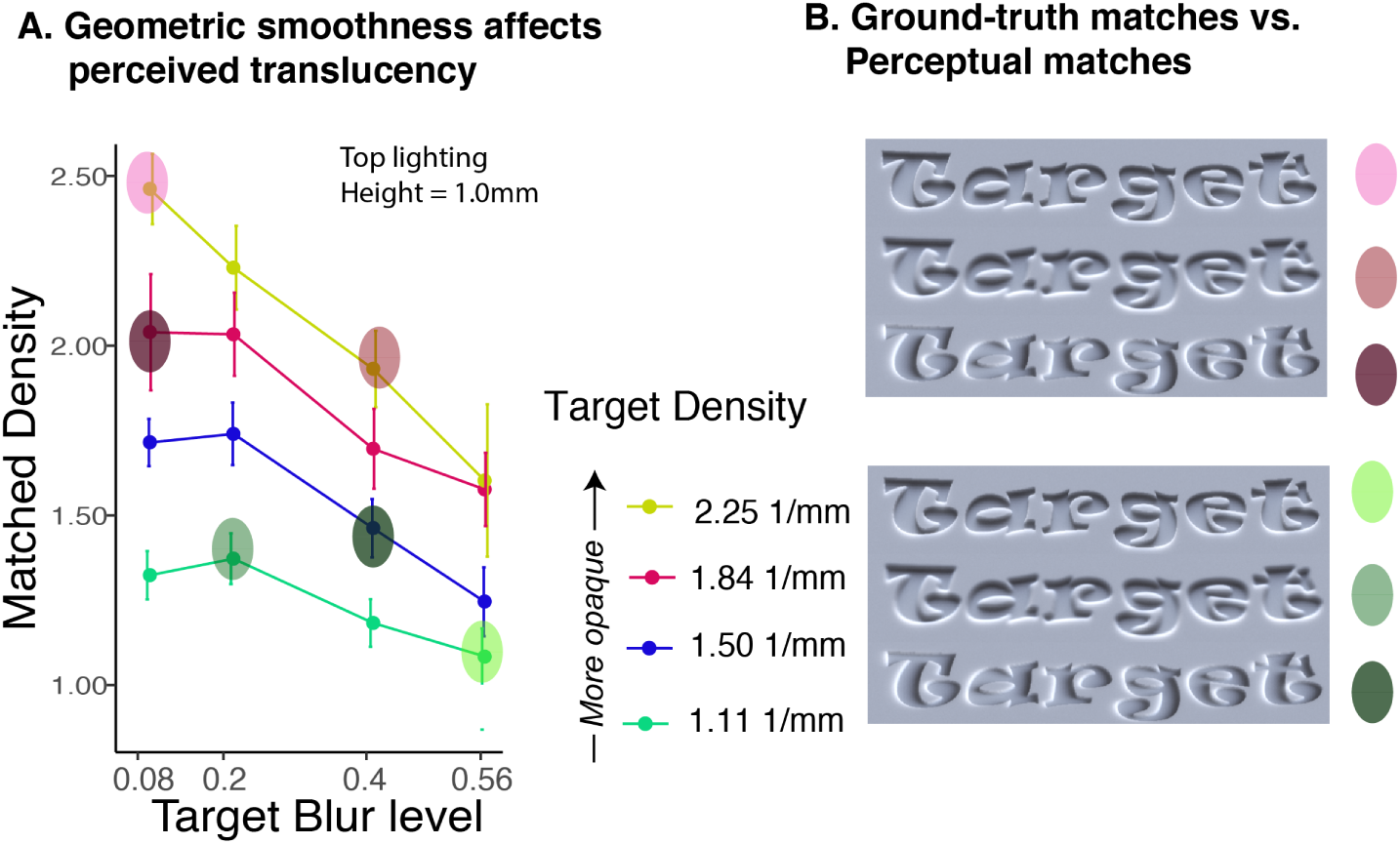
Results from Experiment 2 (Negative relief) of all relief heights (depths) and lightings. **A.** Mean match density versus levels of blur. **B.** Demos of perceptually equivalent image pairs versus pairs that have the same physical densities. Top panel: on average, observers perceive a target object with higher density (middle image, maroon dot) with smooth geometry to be equivalent to an object with lower density but sharp geometry (lower image, darker maroon dot). The top image shows the physical ground-truth of the image of the object with the same density as the target but with a lower blur level (pink). Bottom panel: on average, observers perceive a target object with lower density (middle image, green dot) with sharp geometry to be equivalent to an object with higher density but smooth geometry (lower image, dark green dot). The top image shows the physical ground-truth of the image of the object with the same density as the target (lime dot).

**Figure 12:**
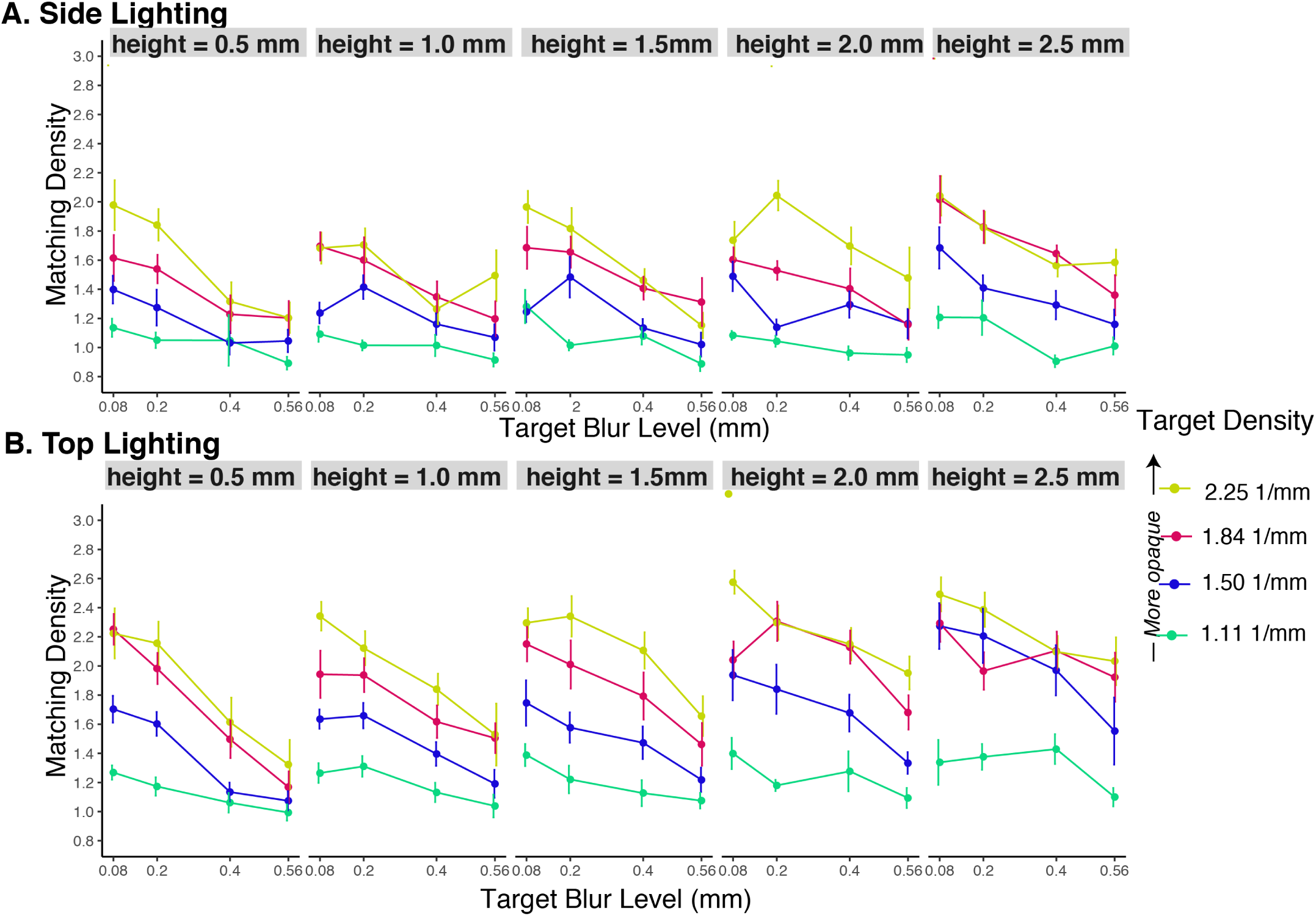
Results from the negative relief: Effect of blurring on the matching results for all relief heights and two lighting directions. Different colors represent different ground-truth densities for the target object. The top panels are data from the side lighting conditions and the bottom panels are data from the top lighting conditions. Similar as the results from Experiment 1, as geometric blur increased for the target, observers’s matched density decreased, suggesting blur affect perceived translucency.

The top panel of Figure 11B shows the effect of blur on translucent appearance of selected images. The middle image of the top panel is a cross section of a blurred object which corresponds to the maroon dot on the data plot in Figure 11A. The bottom image of the panel shows the image rendered with mean matched density across observers (dark maroon). Even though the perceptually matched object in this image has lower density and sharper features (lower blur level) than the target (red dot on data plot in Figure 11A), observers perceive them to be similar. In contrast, the top image shows the object rendered with the same physical density as the target but with a lower blur level (pink dot on the data plot). Observers perceive this image to be more opaque than the target. This illustrate that geometrically blurring the object results in observer perceive the object to be more translucent.

Similar as Experiment 1, bottom triplets of images figure 11 demonstrated how an object with sharp features (middle row, green dot) could be perceived to be more opaque, which is equivalent to the stimuli with higher density (lower panel, dark green dot) than a blurred object that has the same ground-truth density (top panel, lime dot).

Figure 12 shows the mean matched density across observers for the negative relief for all of the conditions. From left to right, the panels show data plots for stimuli with increased relief heights (i.e., deeper relief depths). The two rows show data from the two lighting conditions.

As in Experiment 1, we find that blur has a significant effect on matched density on nearly all height conditions and both lighting conditions. In particular, observers perceive the blurred object to be more translucent than the un-blurred object with the same density. Additionally, and again similar to Experiment 1, blur has a stronger effect for the high density conditions (yellow and red lines) than the low density conditions (dark blue and green lines).

Similar to Experiment 1, we perform a three-way within-subjects ANOVA on the difference between matched density and the target density with blur level, relief height, and lighting direction as independent variables. The results are as follows:

- Similar to Experiment 1, *blur level* has a significant main effect (*F* (3, 1520) = 28.883, *p <* 0.001), such that as the amount of blurring increases, the value of *d*_match_ − *d*_target_ decreases.
- Also similar to Experiment 1, *relief height* also has a significant main effect (*F* (4, 1520) = 7.149, *p <* 0.001), such that higher relief height makes the object appear less translucent (i.e., higher values of *d*_match_ − *d*_target_). There is no significant interaction between blur level and relief height (*F* (12, 1520) = 0.926, *p* = 0.5199).
- Again similar to Experiment 1, *lighting direction* does not have a significant main effect (*F* (1, 1520) = 45.103, *p <* 0.001). There is no significant interaction between lighting and blur level (*F* (3, 1520) = 0.457, *p* = 0.7125).

## 7. Discussion

Our results indicate that the perception of translucency depends not only on an object’s optical parameters (e.g. scattering parameters used in rendering), but also on their 3D shapes. We used an asymmetric task where observers adjusted optical density of a “match” object to match the material properties of a “target”. In Experiment 1 (with positive relief), we find that observers tend to perceive the geometrically smoothed objects as more translucent than those with sharper geometries. This effect is significant across most relief heights and under both side and top lighting conditions except the highest relief under top lighting. In Experiment 2 (with negative relief), we find similar effects of 3D geometry on translucency across all conditions. In the following, we discuss whether observers use low-level image intensity information during the match and whether image contrast can be used to predict the results.

### Predictions from image similarity metrics

To test the hypothesis that observers are not performing low-level image similarities during the match, we first compared a simple least-square image distance (L2 norm) between the “target” and “match”. For each “target” image used in the experiment, we computed the L2 norm between a “target” image and all the “match” images that have the same relief heights and lighting conditions. Then we searched for the matching density that resulted in the smallest L2 distance. In order to correct the effect of mean brightness caused by different optical densities, we normalized both the “target” and the “match” images intensities with their mean luminance values before we compute L2 norm. We used the same set of “match” images human observers used in the matching experiment to compute L2 prediction. Figure 13 shows the results from the predicted matches based on the L2 norm for all “target” conditions used in Experiments 1 and 2. We found the predictions of all target images to be a single value, which is the lowest density 0.74*mm*, which is obviously different from the perceptual results.

**Figure 13:**
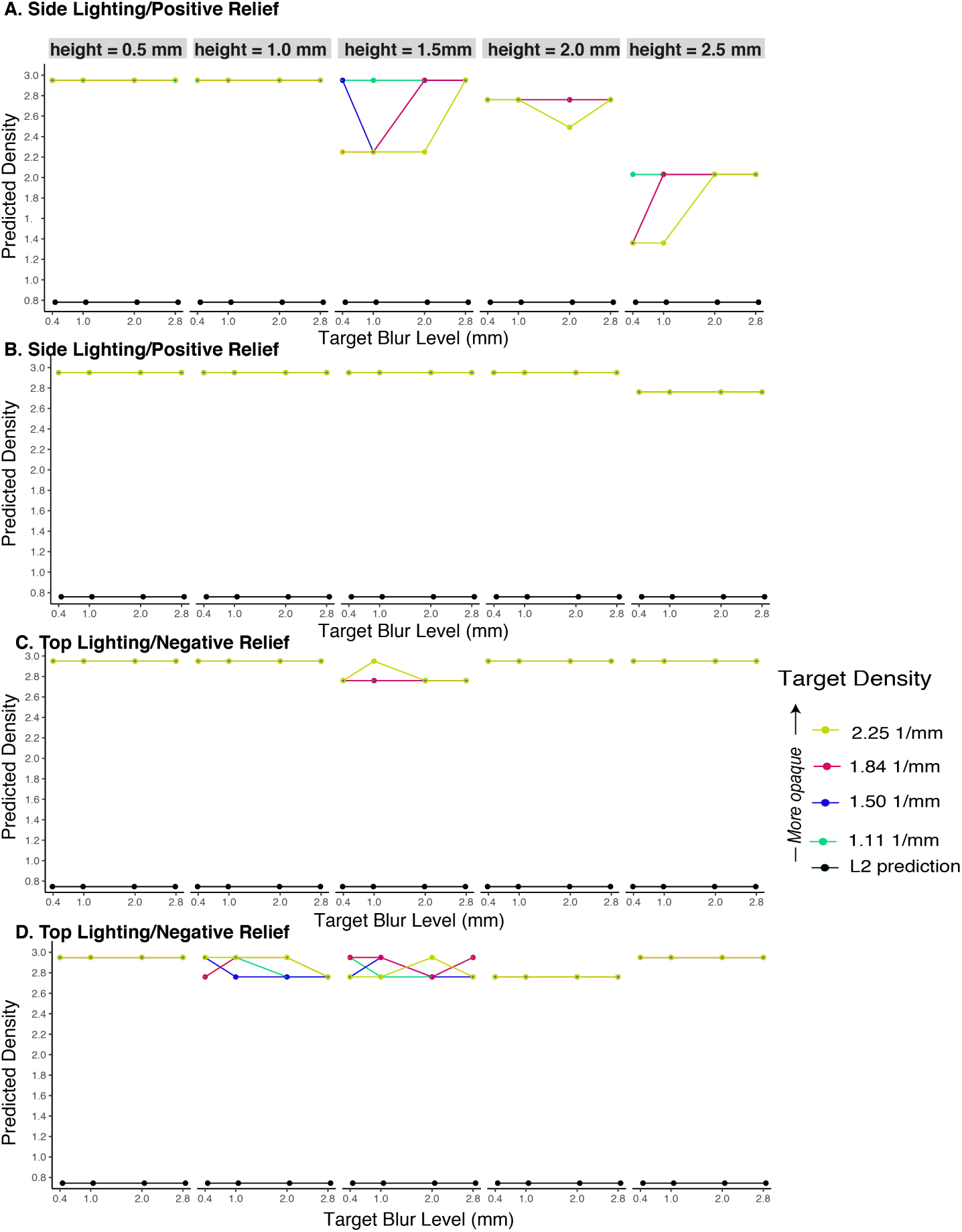
Prediction from image metric: predicted matched density with an image similarity metric (The structural similarity (SSIM) index) and a Euclidean distance metric (L2 norm) for all conditions from both Experiment 1 and 2. The prediction was computed by finding the density that minimizes the SSIM or L2 norm between a “target” image and each of the “match” images observers used in Experiments 1 and 2. The colored dots and lines are prediction from SSIM. Since the predictions from L2 norm collapsed into one value, we used a single black color to plot the predicted value. From top to bottom, the panels plots predictions from the images of both relief and lighting conditions. A. Predictions computed from images of positive relief under “side lighting”. From left to right, the images have increasing relief heights. B. Predictions computed from images of positive relief under “top lighting”. C. Same for images of negative relief under “side lighting”. D. Same for negative relief under “top lighting”. The meaning of the symbols, legends, range of the X and Y axis are the same as those in Figure 9.

L2 norm is known to be a poor indicator of perceptually visible differences between images. Hence, we used a perception-based image similarity metric to compute the predictions. Specifically, we computed structural similarity index (SSIM) (Wang, Bovik, Sheikh, Simoncelli, et al., 2004) between a “target” image and each of the “match” images observers used in Experiments 1 and 2. We then computed the prediction by finding the density that maximize the SSIM. We then can compare the prediction with the perceptual results. Figure 13 shows that the SSIM-based predictions of all target density, blur level, and relief height values used in Experiment 1 and Experiment 2. The figure shows that the prediction from SSIM are rather random and is not at all similar to the perceptual results (see figure 9 and figure 11). In our experiments, we made sure that on an image-level, the “target” image differ in ways other than geometric sharpness than the “match” image such as they are rendered with different relief characters and lighting. Hence, the images are not at all registered and this might have caused both the SSIM and L2 norm being not predictable of the perceived material similarities. In summary, we found both L2 norm and SSIM cannot predict perceptual results, suggesting in our asymmetric matching experiments, observers are unlikely use low-level image cues to make the matches.

### Is image contrast a possible metric for translucency?

We next explore the relationship between image contrast and perceived translucency using our stimuli. Previous work found that the human visual system might use image contrast (Michelson) as a critical image variable to initiate percepts of transparency and to assign transmittance to transparent surfaces (Singh & Anderson, 2002). A further study with more realistic images shows reducing contrast of the background decreased perceived transparency of the overlaying filter (Robilotto & Zaidi, 2004) and studies on volumetric translucency also reveals significant effect of image contrast (Fleming & Bülthoff, 2005; Motoyoshi, 2010). Here, we computed the Michelson contrast between “target” and the candidates “match” images ((*I*_max_ − *I*_min_)*/*(*I*_max_ + *I*_min_)) and use the contrast to make predictions of densities in a similar way as what has bee described in above section.

Figure14 plots the predicted density based on Michelson contrast between “target” and “match” images for all experimental conditions. First of all, image contrast could discriminate different optical densities as good as the perceptual data. Second, the effect of blur resembles perceptual data for the shallow relief such that as blur increases, the predicted density decreases (see left two columns in figure 14. However, as the relief height increases, the effect of blur on predicted density is diminished if not reversed (see right two columns in figure 14 in bottom raws), whereas in perceptual data, increasing blur would cause matched density to decrease across all reliefs (see figure 9 and figure 11). Hence, the predictions generated from image-contrast cannot fully predict human perception data, suggesting that even though image-contrast could be useful for some conditions, it is certainly not the only image cue the visual system used in our experiments.

**Figure 14:**
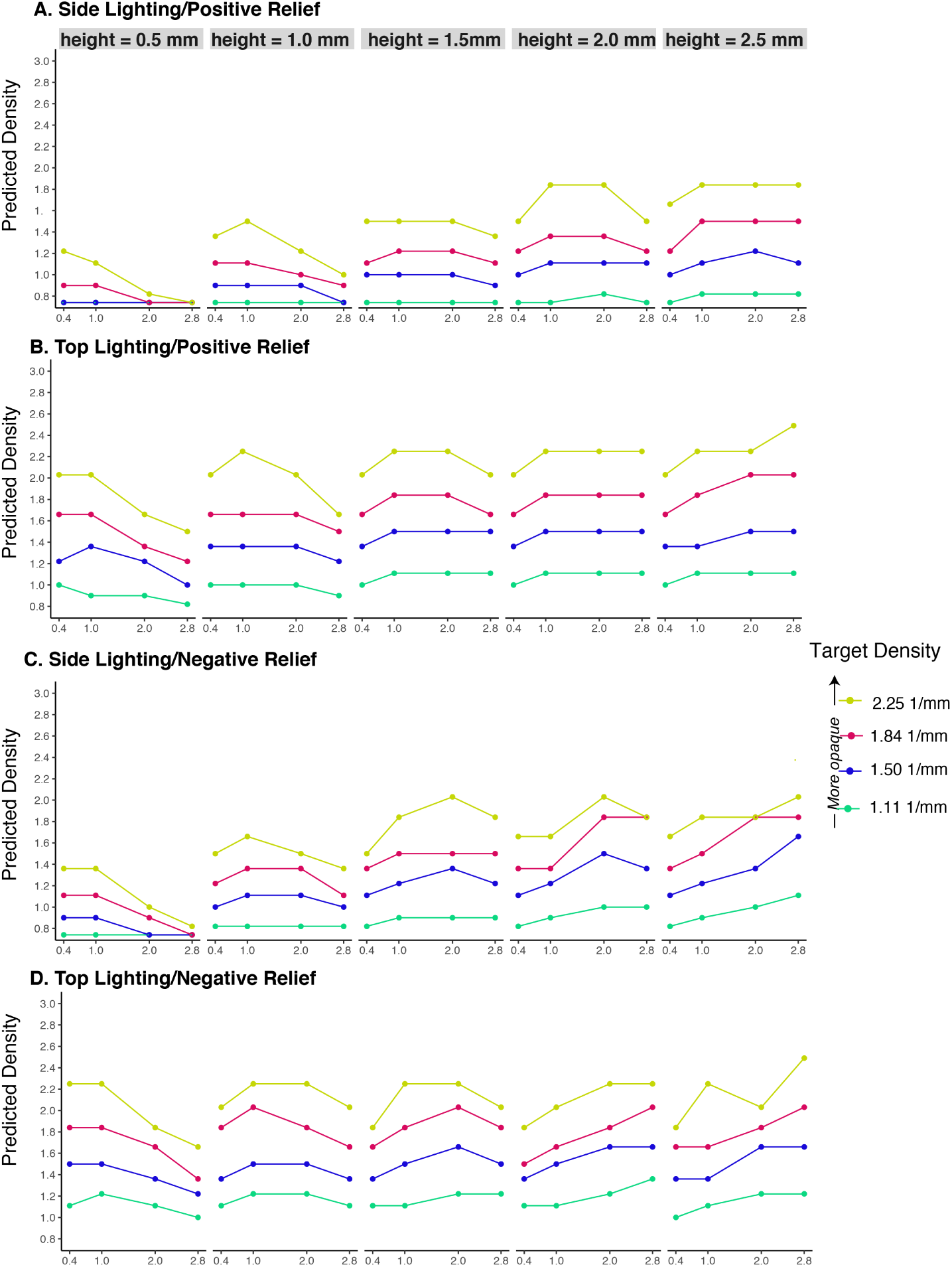
Prediction from image contrast: predicted matched densities from comparing Michelson Contrast (see texts for details) between a”target” image and each of the candidate “match” images for all conditions in Experiments 1 and2. The panels are organized in the same way as figure 13. The meaning of the symbols, legends, range of the X and Y axis are the same as those in Figure 9.

## 8. Conclusion

We performed psychophysical experiments, using physically based rendering to synthesize visual stimuli, and asked observers to match the perceived level of translucency of stimuli showing objects of different 3D geometric sharpness. We discovered that the perceived translucency depends not only on the material scattering parameters, but also on the objects’ 3D geometries. Observers tend to perceive geometrically smooth objects to be more translucent than geometrically sharp objects, across different lighting direction and relief shape. We also found that simple image-based metrics (L2-distance and structural similarity index) could not predict our perceptual data, further suggesting observers were not use intensity-based image similarity to make the judgements. Analysis of Michelson contrasts of the images suggests even though image contrast could be a cue for estimating translucency for some shape conditions, it cannot fully predict our perceptual data. This suggests contrast is not the only cue used by the visual system. Our results indicate that modifying the finely detailed 3D shapes of translucent objects could also alter their perceived appearance.

The findings reported in this paper have implications in designing perception-aware metrics for translucency and 3D fabrication. Translucent objects with the same material properties but different fine 3D shape might appear different. In contrast, it is also possible to fabricate objects with the same overall perceived appearance, by using different physical materials and carefully manipulating the objects’ fine 3D geometry. This suggests that, when fabricating objects with an objective to match a specific material appearance, their 3D shape must also be considered.

## Footnotes

1 We provided the examples of tonemapped stimuli images in the supplementary material. The full datasets, individual subjects’ data, and the user-interface code will be available upon acceptance of our paper.

